## Supplementary figures 1-5 for "Subtypes and proliferation patterns of small intestine neuroendocrine tumors revealed by single cell RNA sequencing"

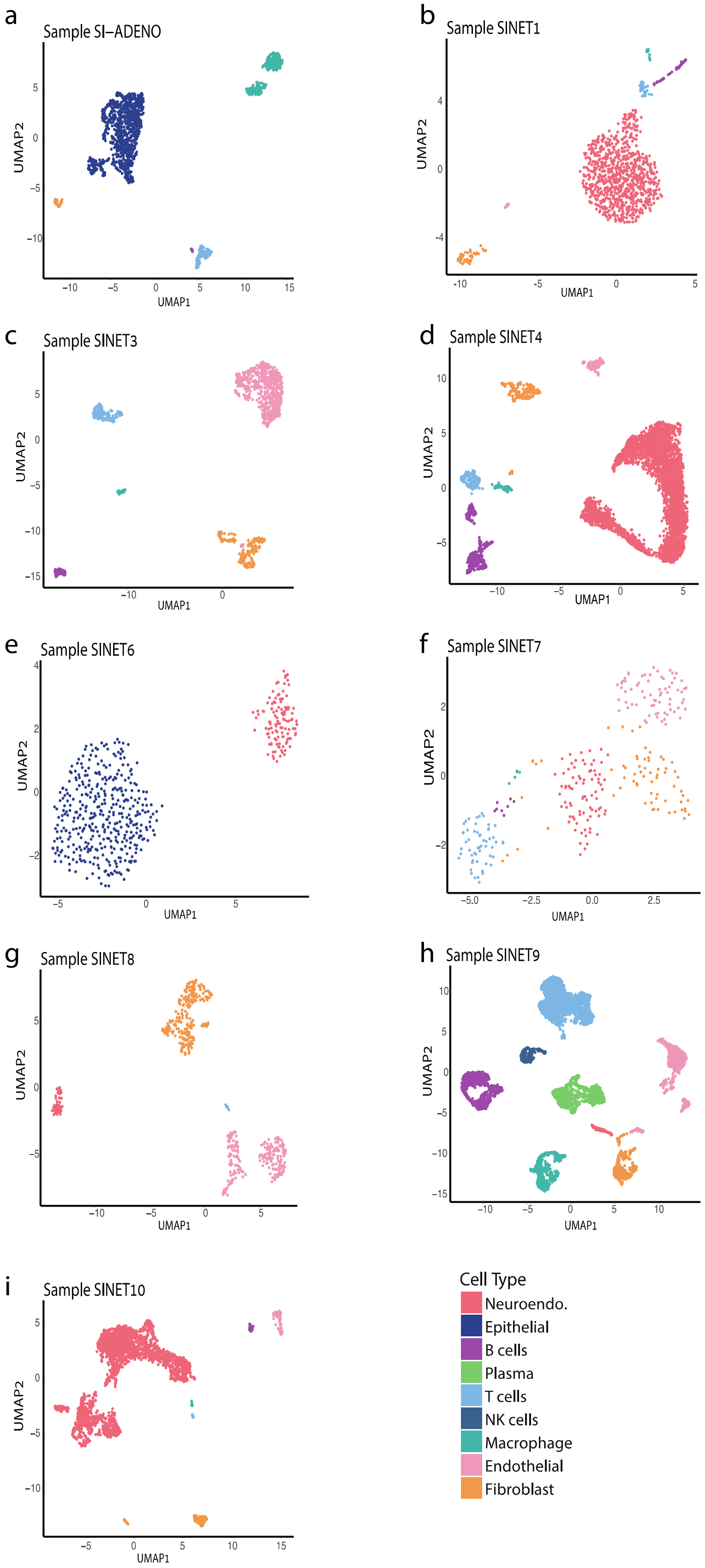


**Figure S1. Cellular composition of SiNETs as determined by single cell and single nuclei sequencing.** (**A-I**) UMAP plots showing the diversity of single cells from each sample, colored by their clustering.

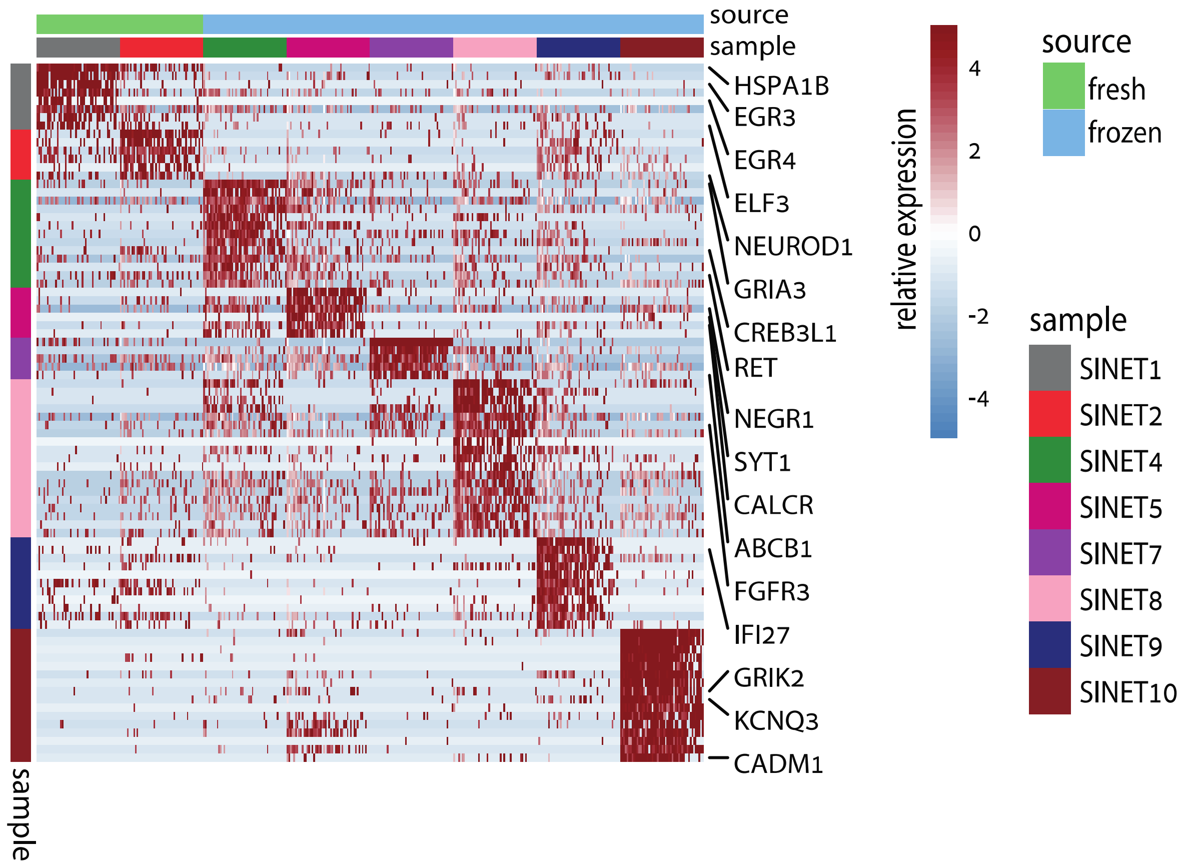


**Figure S2. Specific upregulated genes expressed in neuroendocrine cells per siNET sample.**

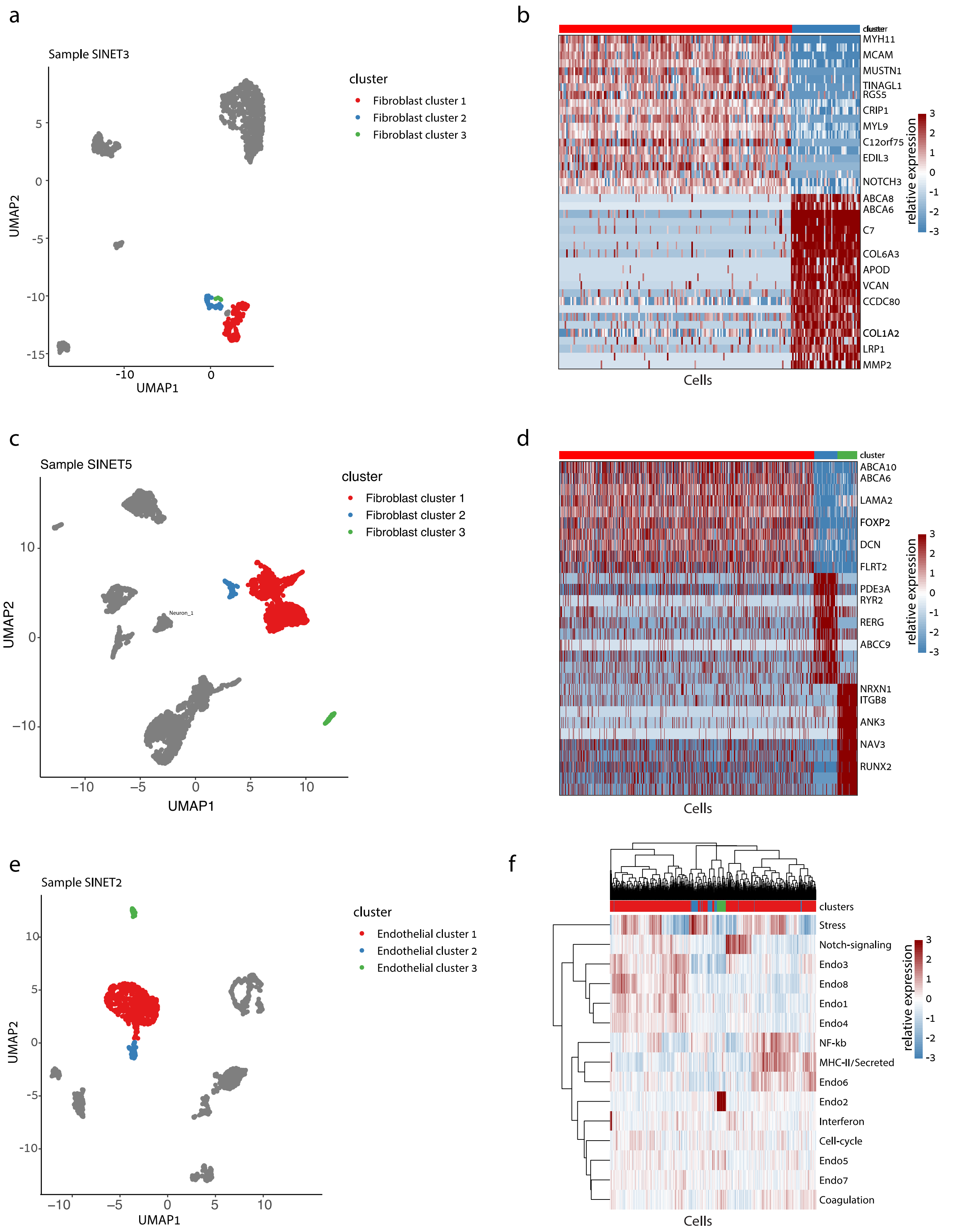


**Figure S3. Heterogeneity in the SiNET tumor microenvironment.** For each of three non-malignant cell types, the diversity of that cell type is shown in one exemplary tumor: fibroblast heterogeneity in SiNET3 is shown in (**A, B**), fibroblast cell heterogeneity in SiNET5 is shown in (**C, D**), and endothelial heterogeneity in SiNET2 is shown in (**E, F**). For each cell type, panels depict types of analyses. The first panel (**A, C, E**) is a UMAP plot of the respective tumor, where only the respective cell type is colored, and distinct colors highlight the clusters of that cell type. The second panel (**B, D**) shows differential expression analysis between the first two clusters using a heatmap, with labeling of selected genes. The third panel (**F**) shows clustering of cells from that cell type (columns) based on their relative expression of previously defined [18] signatures of diversity in that cell type (rows); the top panel shows assignment of cells to clusters.

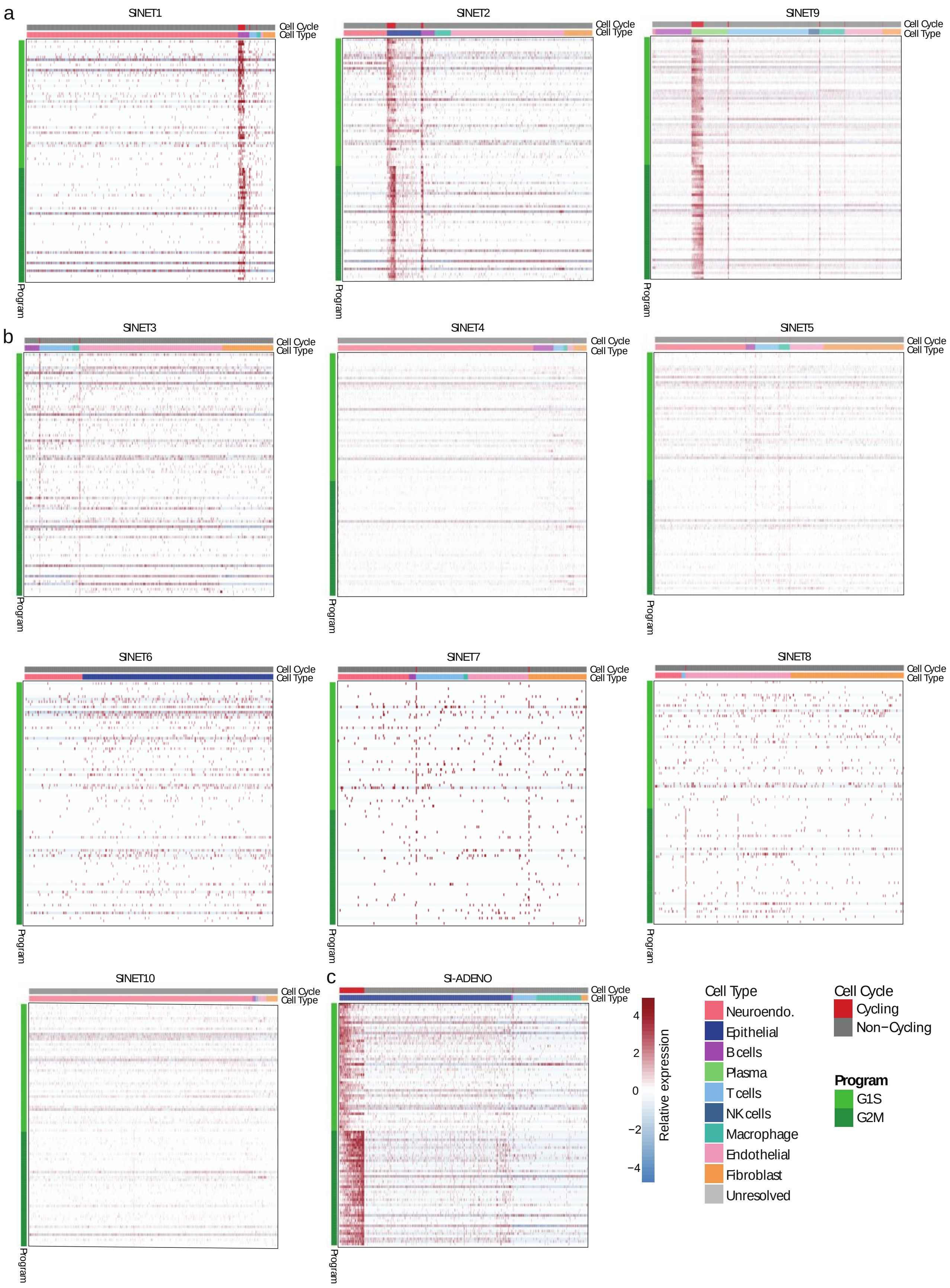


**Figure S4. Heat map illustrating the expression of G1/S and G2/M genes across various cell types in the Epithelial-like Subtype (A), Neuronal-like Subtype (B) and the siAdeno sample (C).** The annotated cell types include Epithelial cells, Macrophages, T cells, and Fibroblasts that were sampled for illustrative purposes.

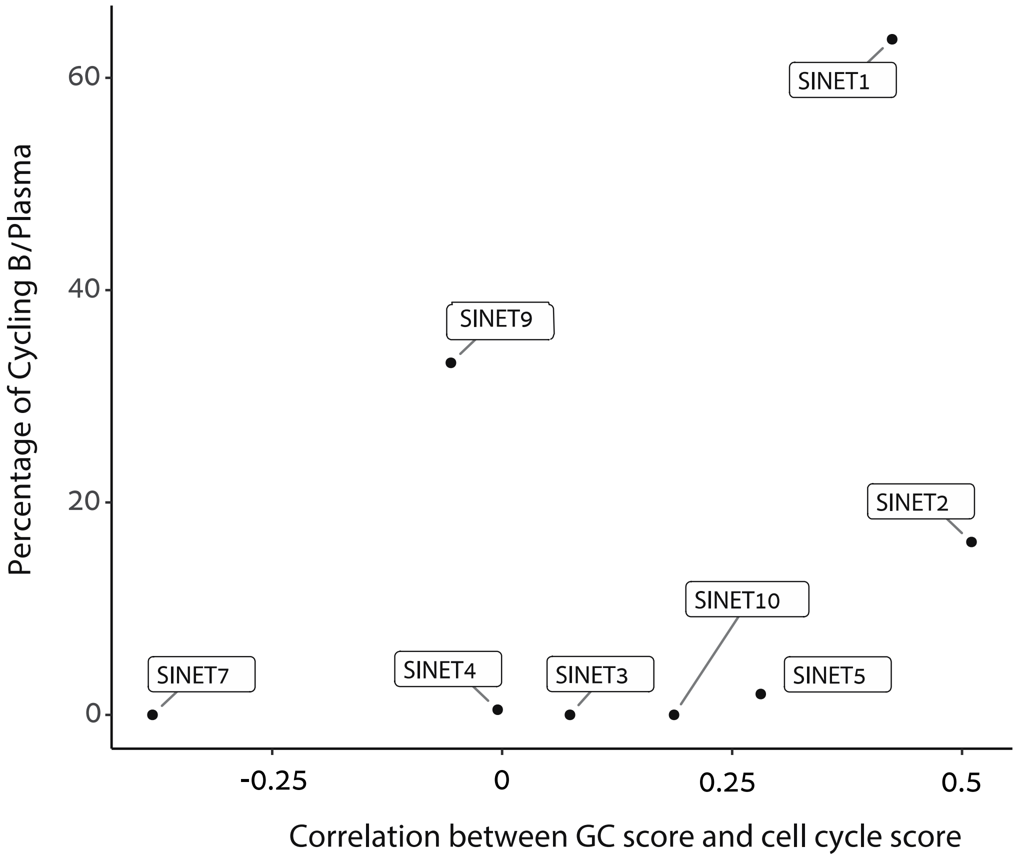


**Figure S5. Scatter plot illustrating the percentage of cycling B/Plasma cells and the correlation between the germinal center signature and cycling B/Plasma cells signature.** In SiNET1 and SiNET2 we observe high correlation between cell cycle and GC score, but not in SiNET9.
